## Supplemental Information for "Dynamics of nevus development implicate cell cooperation in the growth arrest of transformed melanocytes"

**Supplemental Material**

**Data S1 (uploaded as .xlsx)**

**Data S2 below:**

**Data S2. Parameters values used in CompuCell3D model of nevus growth arrest (**refers to Figure 5C-D).

| **Property** | **Value** | **Units** | **Comments** |
|---|---|---|---|
| *Simulation Domain* | | | |
| Monte Carlo Steps in simulation | 30000 | | |
| Lattice Width | 300 | px | Note 1 |
| Lattice Area | 900 | px^2 | |
| *Cell proliferation* | | | |
| Cell Target Area | 36 | px^2 | |
| growth | 0.05, 0.1, 0.15 | | Note 2 |
| mitosis | 72-80 | px^2 | Note 3 |
| *Contact Energy* | | | |
| Medium-Medium | 0 | J | |
| Medium-Dividing | 10 | J | |
| Medium-Non dividing | 5 | J | |
| Dividing-Dividing | 15 | J | |
| Dividing-Non dividing | 15 | J | |
| Non dividing-Non dividing | 2 | J | |
| *Diffusible Signal* | | | |
| Secretion | 0.0025 | MCS^-1 | |
| Diffusion Constant | 6 | px^2/MCS | |
| Decay Constant | 0.026667 | MCS^-1 | |
| Transisiton (differentiation) probability | $\alpha/(1+\alpha)$ | | Note 4 |

Note 1 Abbreviations: px = pixel, MCS = Monte Carlo Step

Note 2 *random.choice():* after every cell division choose between these three values

Note 3 *random.choice(range()):* after every cell division choose an integer between 72 and 80.

Note 4 $\alpha$ = the concentration of the diffusible molecule at the center of mass of the cell, divided by 0.3

### Supplemental Figures S1-S4

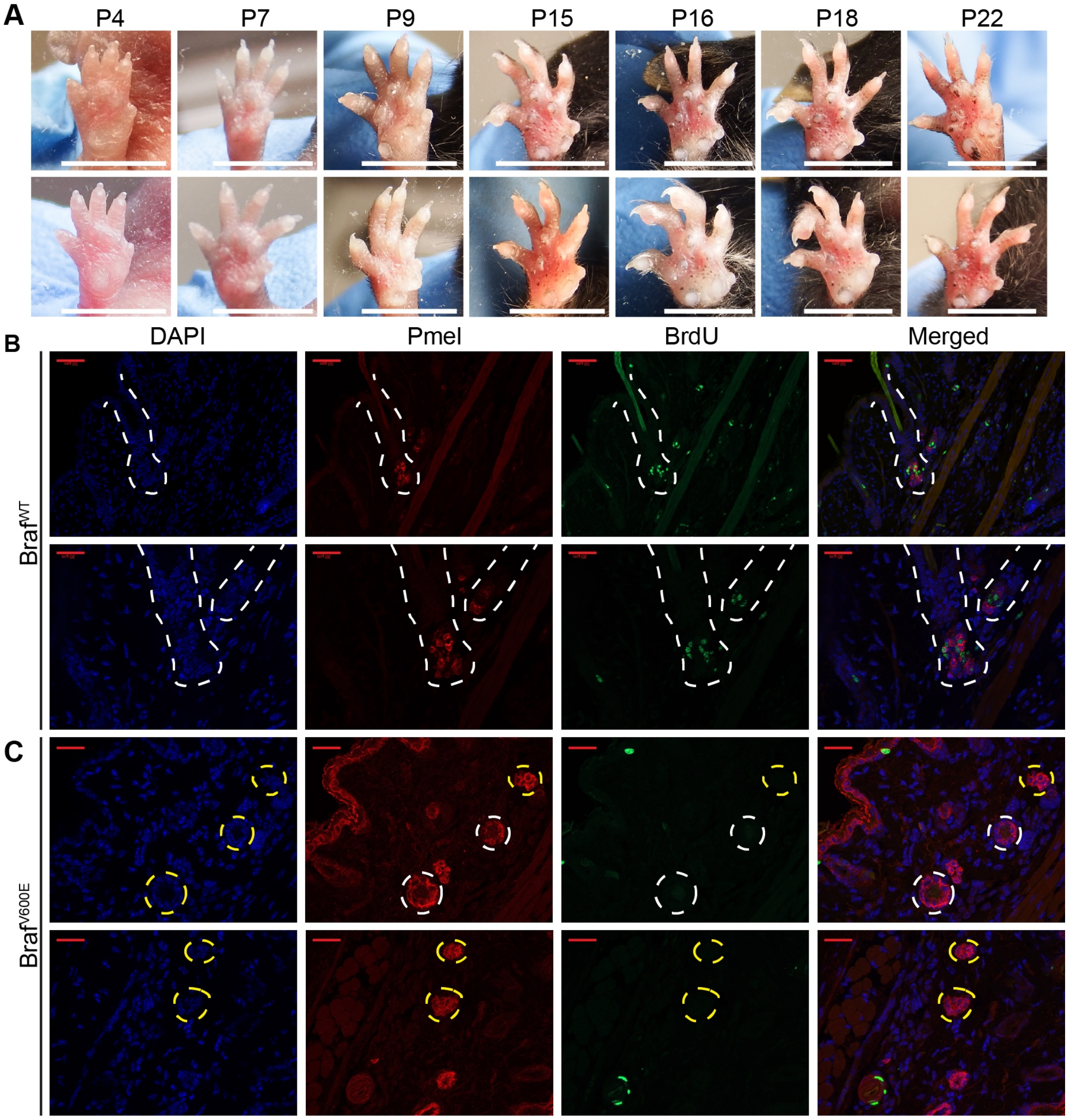

**Figure S1. Dynamics of nevus growth. A.** Nevus development on the ventral surface of the glabrous paw. Images were taken at the indicated age. **B-C.** Identification of dividing or non-dividing melanocytes with the premelanosome protein (Pmel) and BrdU incorporation in the **B.** hair follicle (HF) of $Braf^{WT}$ and **C.** in nests at P21 in $Braf^{V600E}$ mice. Scale bar = 20μm.

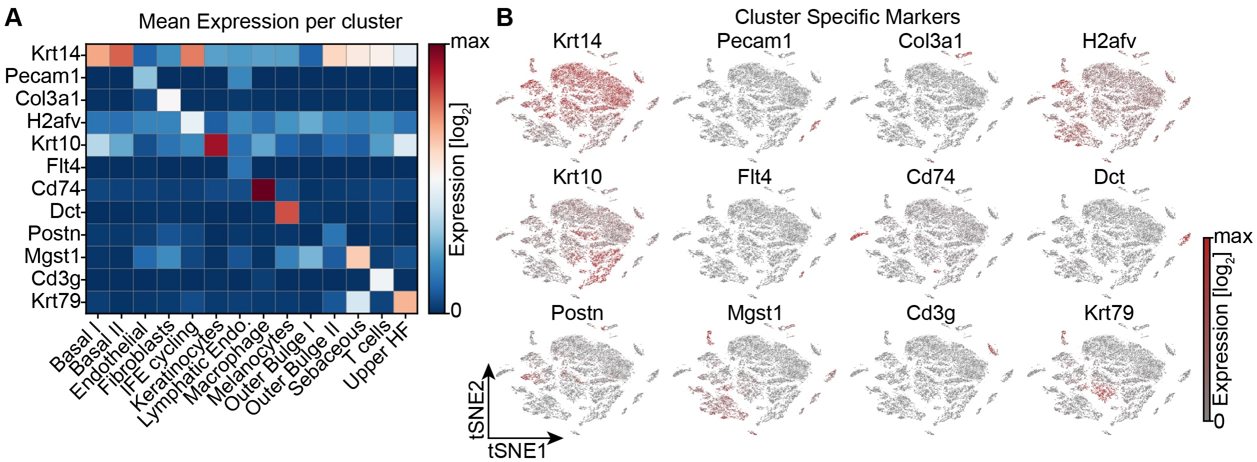

**Figure S2. Single cell RNA sequencing of mouse dorsal skin to transcriptionally characterize cells. A.** Heat map depicting the average gene expression of canonical markers for known cell types found in the skin. **B.** Gene expression of the indicated gene in cells to identify cell type clusters.

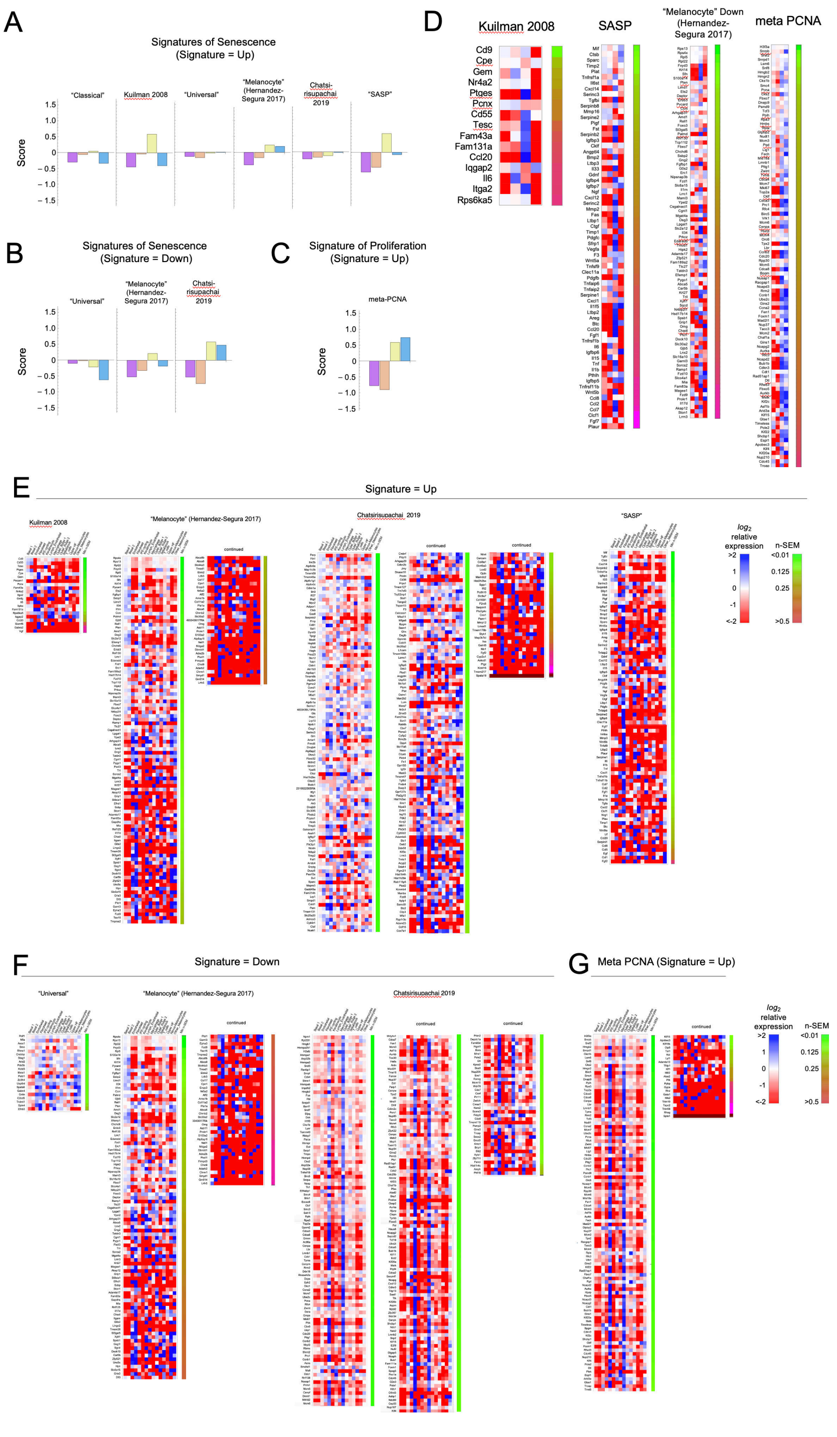

**Figure S3. Heat maps for other signatures associated with senescence or proliferation.** The single cell transcriptomes analyzed in Figure 3 were compared with each of the signatures in Data table S1. **A-D.** Summary (A-C) and detailed (D) comparisons for the four melanocytes subclusters. **E-F**. detailed comparisons of all skin subclusters to all signatures not shown in Fig. 3.

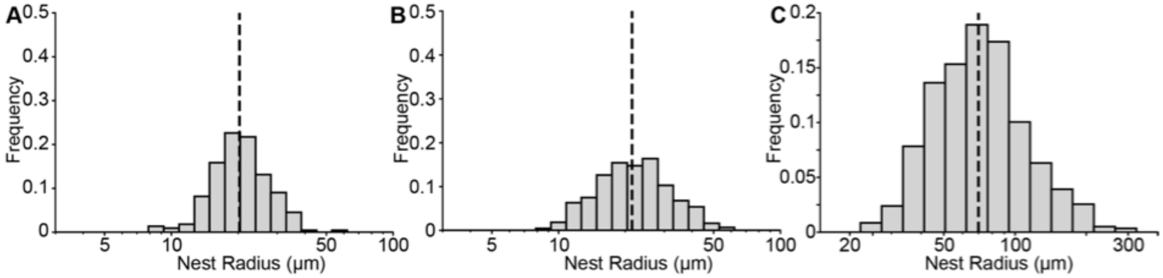

**Figure S4. Size distribution of nests in both mouse and humans. A-B.** Quantification of nest radii treated with 75mg/mL of tamoxifen at **(A)** P21 (mice = 10, nests = 221) or **(B)** P50 (mice = 18 mice, nests = 428). The black dashed line represents the median of 20.2μm and 21.4μm, respectively. **C.** Quantification of human nests. The black dashed line represents the median radius of 70μm.

### Mathematical Supplement to "Dynamics of nevus development implicates cell cooperation in the growth arrest of transformed melanocytes." by Ruiz-Vega et al.

#### Modeling Oncogene Induced Senescence as a Cell Autonomous Process

Consider the clonal descendants of an oncogene-transformed cell. The simplest model of a cell-autonomous oncogene-induced arrest process is one in which cells replicate at a constant rate, and undergo senescence with a constant probability *s* per cell cycle. Once all the cells in a clone have undergone senescence, we refer to the clone as "extinct". For any given *s*, we wish to find the probability that extinction has occurred by any given time, as well as the distribution of clone sizes that should be expected at that time.

If the decision to senesce is independent in each daughter cell at each cell division, then this scenario describes a branching process in which each cell produces two senescent cells with probability $s^2$, two dividing cells with probability $(1\text{-}s)^2$, and one dividing and one senescent cell with probability $2s(1\text{-}s)$. The theory of branching processes may then be used to obtain the probability generating function (PGF) for the number of dividing cells after *n* cell cycles. The offspring distribution of a single cell as described above may be written as:

$$f(z) = s^2 + 2s(1\text{-}s)z + (1\text{-}s)^2 z^2$$

where *z* is a dummy variable whose exponent indicates the number of dividing (non-senescent) cells produced by an event, and the coefficient in front of each $z^n$ is the probability of that event. It can be shown that the PGF for the number of dividing cells after *n* cell cycles is equal to the *n*-th composition of the offspring distribution *f* onto itself (*1*). For example, the PGF for the number of dividing cells after 2 cell cycles is:

$$F(2,z) = f(f(z)) = s^2 + 2s(1\text{-}s)(s^2 + 2s(1\text{-}s)z + (1\text{-}s)^2 z^2) + (1\text{-}s)^2(s^2 + 2s(1\text{-}s)z + (1\text{-}s)^2 z^2)^2$$

And in general:

$$F(n, z) = F(1, F(n\text{-}1, z)) = f(F(n\text{-}1, z))$$

Thus $F(n, 0)$ is the cumulative probability that there are 0 dividing cells—i.e. a clone has extinguished—after *n* cell cycles.

The theory of branching processes also tells us that the probability that a clone eventually goes extinct (over the long run) is just the smallest non-negative root of $z = f(z)$. We solve the equation: $z = s^2 + 2s(1\text{-}s)z + (1\text{-}s)^2z^2$ and obtain: $z = s^2/(1\text{-}s)^2$, $0<s<0.5$ and $z=1$, $s>0.5$. This confirms that only if $s>0.5$ is eventual extinction of all clones guaranteed.

#### Monte Carlo Simulations

The behavior of the above branching processes may be easily observed using Monte Carlo Simulation, seeding each clone with a single dividing cell and specifying the value of *s*. In the simulations show in Figure 4, cells divided synchronously and *s* was used to determine the fate of each daughter cell after each division. For each simulation it was recorded (1) whether the clone went extinct, and if so, after how many cell cycles it did so and (2) the number of cells at the time of extinction (of the end of the simulation).

#### Distribution of clone sizes at extinction

To derive the distribution of clone sizes at extinction it is helpful to think about clonal development not in terms of the number *n* of cell cycle times that have elapsed, but in terms of the cumulative number of cell division events that have occurred up to any given time within a clone, which we will represent as $\eta$. Assuming clones begin from one cell, the number of cells at extinction ($T_c$) will simply be $T_c = 1 + \eta$;

To find the expected distribution of values of $T_c$, we begin by computing the probability that a clone goes extinct at a given value of $\eta$, which we will call $p_E(\eta)$. Recalling that two choices are made at every division (one per daughter cell), it may be seen that to extinguish at $\eta$ requires 2 $\eta$ choices, exactly $\eta$ +1 of which are choices to become senescent, and $\eta$ -1 events are choices to remain proliferative. This implies that:

$p_E(1) = A_1\, s^2$
$p_E(2) = A_2\, (1\text{-}s)s^3$
$p_E(3) = A_3\, (1\text{-}s)^2 s^4$
...
$p_E(\eta) = A_\eta\, (1\text{-}s)^{\eta-1} s^{\eta+1}$

where the coefficients $A_\eta$ are constants that capture the number of different ways that each combination of choices of *s* and 1-*s* can happen. $A_\eta$ can be thought of as the number of unique full binary trees that end with $\eta$ + 1 senescent cells. As senescent cells are the dead ends in those trees, they are referred to as "leaves" of the tree. The sequence that counts the number of unique full binary trees is called the *Catalan numbers*. $C_k$, the *k*th Catalan number is the number of unique binary trees with *k*+1 leaves. It is defined by:

$$C_k = \frac{1}{k+1}\binom{2k}{k}$$

Accordingly,

$$p_E(\eta) = \frac{1}{\eta+1}\binom{2\eta}{\eta}(1-s)^{\eta-1}s^{\eta+1}$$

Consequently, the probability of $T_c$ = *m* cells at extinction is simply

$$p_{T_c}(m) = p_E(m-1)$$

Below, at left, we plot the value of $p_E(\eta)$ as a function of $\eta$ for different values of *s*, using logarithmic axes. Notice that for *s* sufficiently close to 0.5, the relationship approximates a line of slope -3/2. Thus, the approximate probability of finding a clone of size *m* varies inversely with the 3/2-power of *m*. At right, the analytical results for *s*=0.56 are superimposed on results obtained by Monte Carlo simulation of 500,000 cases.

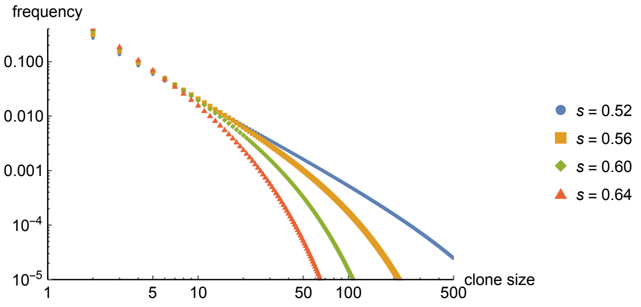

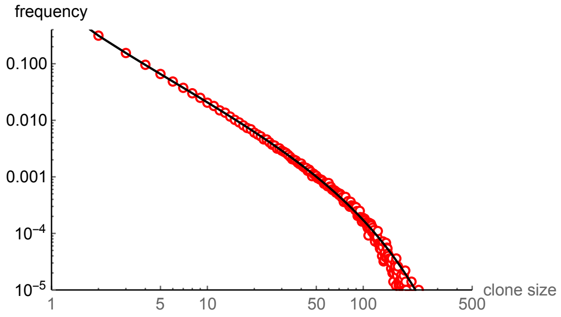

*1*. Hawkins, D., and S. Ulam. "Theory of multiplicative processes I." Los Alamos Scientific Laboratory, LADS-265 (1944).
